# Regionalization, constraints, and the ancestral ossification patterns in the vertebral column of amniotes

**DOI:** 10.1101/2021.09.23.461462

**Authors:** Antoine Verrière, Nadia B. Fröbisch, Jörg Fröbisch

## Abstract

The development of the vertebral column has been studied extensively in modern amniotes, yet many aspects of its evolutionary history remain enigmatic. Here we expand the existing data on four major vertebral developmental patterns in amniotes based on exceptionally well-preserved specimens of the early Permian mesosaurid reptile *Stereosternum:* (i) centrum ossification, (ii) neural arch ossification, (iii) neural arch fusion, and (iv) neurocentral fusion. We retrace the evolutionary history of each pattern and reconstruct the ancestral condition in amniotes. Despite 300 million years of evolutionary history, vertebral development patterns show a surprisingly stability in amniotes since their common ancestor. We propose that this conservatism may be linked to constraints posed by underlying developmental processes across amniotes. However, we also point out that mammals and birds differ more strongly from the ancestral condition than other clades, which might be linked to a stronger regionalization of the column in these two clades.

## Introduction

The vertebral column is the defining feature of all vertebrates and has long been the subject of special attention by developmental and evolutionary biologists, yet certain key aspects of its development and evolution remain unexplored. Thanks to a large body of research spanning more than a century (Gadow, 1896; Kölliker, 1876; Lawson and Harfe, 2017), the way vertebrae are formed throughout ontogeny is generally well understood. Vertebrae ossify via endochondral ossification, meaning that embryonic cartilaginous frameworks progressively mineralize into fully ossified elements. Ossification starts at one or several centers of ossification inside the cartilage matrix and spreads from there until the matrix is entirely replaced by bone.

In amniotes, elements of the vertebral column undergo two stages of endochondral ossification. The first stage, referred to as primary ossification, consists of the mineralization of the neural arch and pleurocentrum. As the amphibian intercentrum is lost in amniotes (Clack, 2012), we only consider the pleurocentrum here and refer to it as centrum for the purpose of this study. The paired elements of the neural arch arise from two ossification centers, one on each side, and eventually fuse dorsomedially. The mineralization of the centrum starts in a pair of ventrally located ossification centers (Danto et al., 2017) and then spreads through the centrum. Eventually, centrum and neural arch come into contact forming what is called the neurocentral suture. It is only much later in development that this suture closes and both elements merge into a single vertebral unit in most amniotes. The second stage of vertebral ossification, or secondary ossification, is marked by the mineralization of the transverse processes and the spinous process.

Ossification events do not occur simultaneously throughout the vertebral column. On the contrary, they can originate in different locations along the vertebral column, occur at different times, and progress at different speeds and in different directions. In a given ossification pattern, it is generally possible to identify one or multiple spots from which ossification spreads, whether it is the first vertebrae where ossification centers are visible or the first vertebrae in which given elements begin to fuse. For clarity, we will indistinctly refer to these pattern starting points for initial ossification as loci (Fig. 1). Loci are not to be confused with ossification centers, which are the points where ossification begins within a single vertebra.

**Figure 1.**
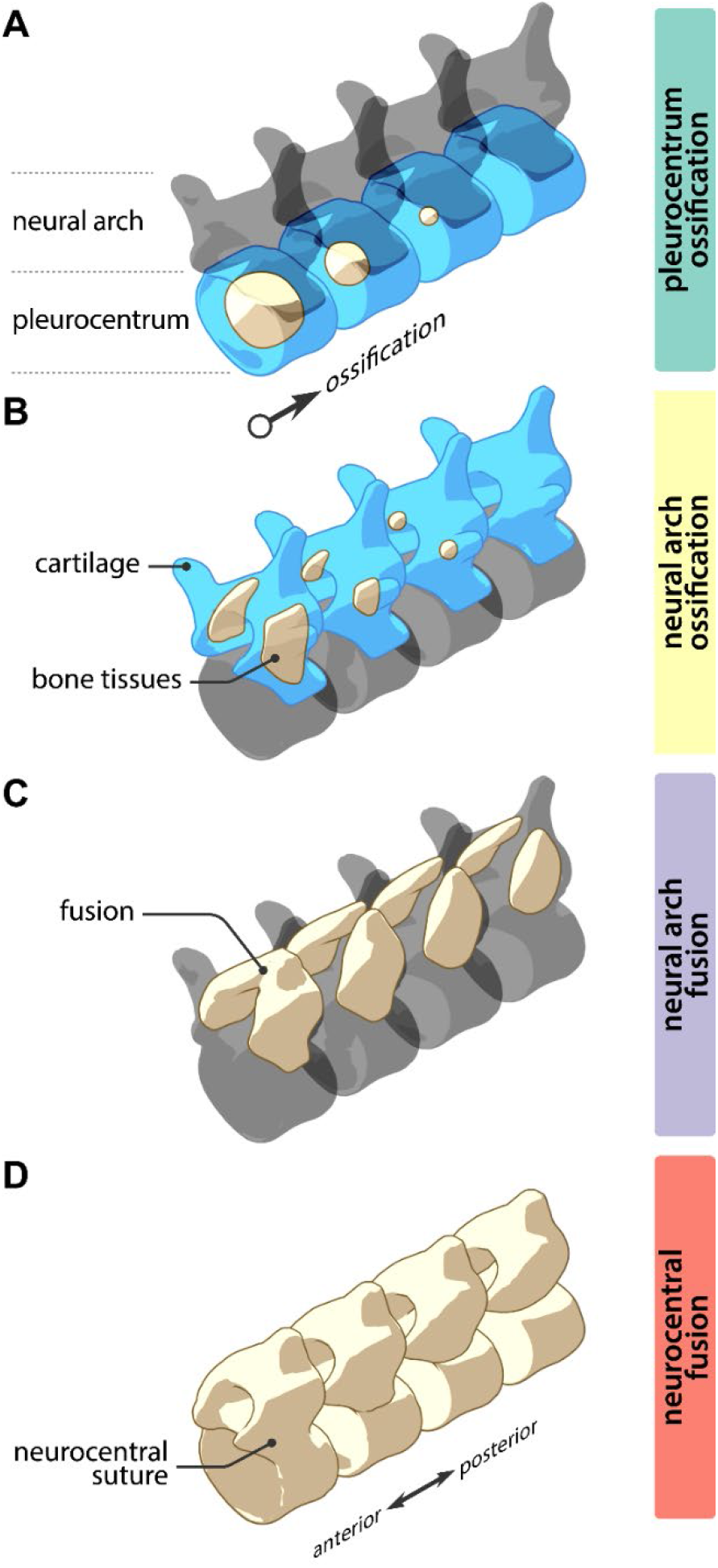
Schematic illustration of the four vertebral ossification patterns examined in the present study. A. pleurocentrum ossification. B. neural arch ossification. C. neural arch fusion. D. neurocentral fusion.

Although the sequence of ossification events leading to the formation of a single vertebra in amniotes has been extensively studied (Böhmer, 2017; Bui and Larsson, 2021; Zhang, 2009), the timing of occurrence and spatial progression of these events along the vertebral column has rarely been documented.. Even for model organisms, most studies only mention which of the neural arches or centra ossify first, if at all. With few exceptions (Hautier et al., 2011, 2010), the disparity between patterns of vertebral development in amniotes remains virtually unstudied, especially with respect to their evolutionary history. While this is partly due to the infrequent preservation of ontogenetic series in the fossil record, the axial column has also suffered from a limited research interest in developmental paleontology. Neurocentral fusion (fusion of neural arches to the centra) constitutes the only notable exception to this as it has been used as a proxy for maturity in fossil archosaurs, especially in dinosaurs. Consequently, this fusion pattern is relatively well documented for fossil members of this clade, albeit in a different context than in this study (Irmis, 2007).

Vertebral elements are considered serially homologous within Amniota (Fleming et al., 2015), which allows for comparisons between clades. Here, we review the current state of knowledge on the four major patterns of primary ossification of the vertebral column in living and fossil amniotes (Fig. 1): (i) the ossification of pleurocentra (ii) the ossification of neural arches, (iii) the fusion of paired neural arch elements, and (iv) neurocentral fusion. These patterns are easily observed on the skeleton and therefore more likely to be preserved in a fossil. In addition, thanks to some exceptionally well-preserved fossils of the early Permian mesosaurid reptile *Stereosternum tumidum,* we provide the first documentation of vertebral ossification patterns in an early amniote. We compiled data on each of these patterns from the literature and used ancestral state reconstruction (ASR) to trace their evolutionary history in the amniote clade. Finally, we reconstructed the hypothetical ancestral condition for each of the patterns in amniotes.

Remarkably, we find that despite the 300-million-year history and great morphological diversity, axial ossification patterns seem to be relatively conservative within Amniota. Our results also show that the evolution of axial development was marked by the additional acquisition and shifting of loci in several clades of amniotes, notably in mammals and birds, which may be linked to the increased regionalization of the vertebral column in these two clades.

## Results

### Centrum ossification

We were able to document the primary sequences of vertebral ossification in the fossil mesosaurid reptile *Stereosternum* based on the juvenile specimen SMF-R-4512. In this specimen, we could identify differences in the degree of ossification of some bones based on variations in color and robustness. SMF-R-4512 shows a stronger coloration and thicker bone in the centra and neural arches of the cervical vertebrae as compared to other regions of the vertebral column (Fig. 2A), indicating a more advanced ossification in this region. This suggests that, in *Stereosternum,* centra and neural arches each had a single anterior cervical locus of ossification and that ossification progressed posteriorly from there.

**Figure 2.**
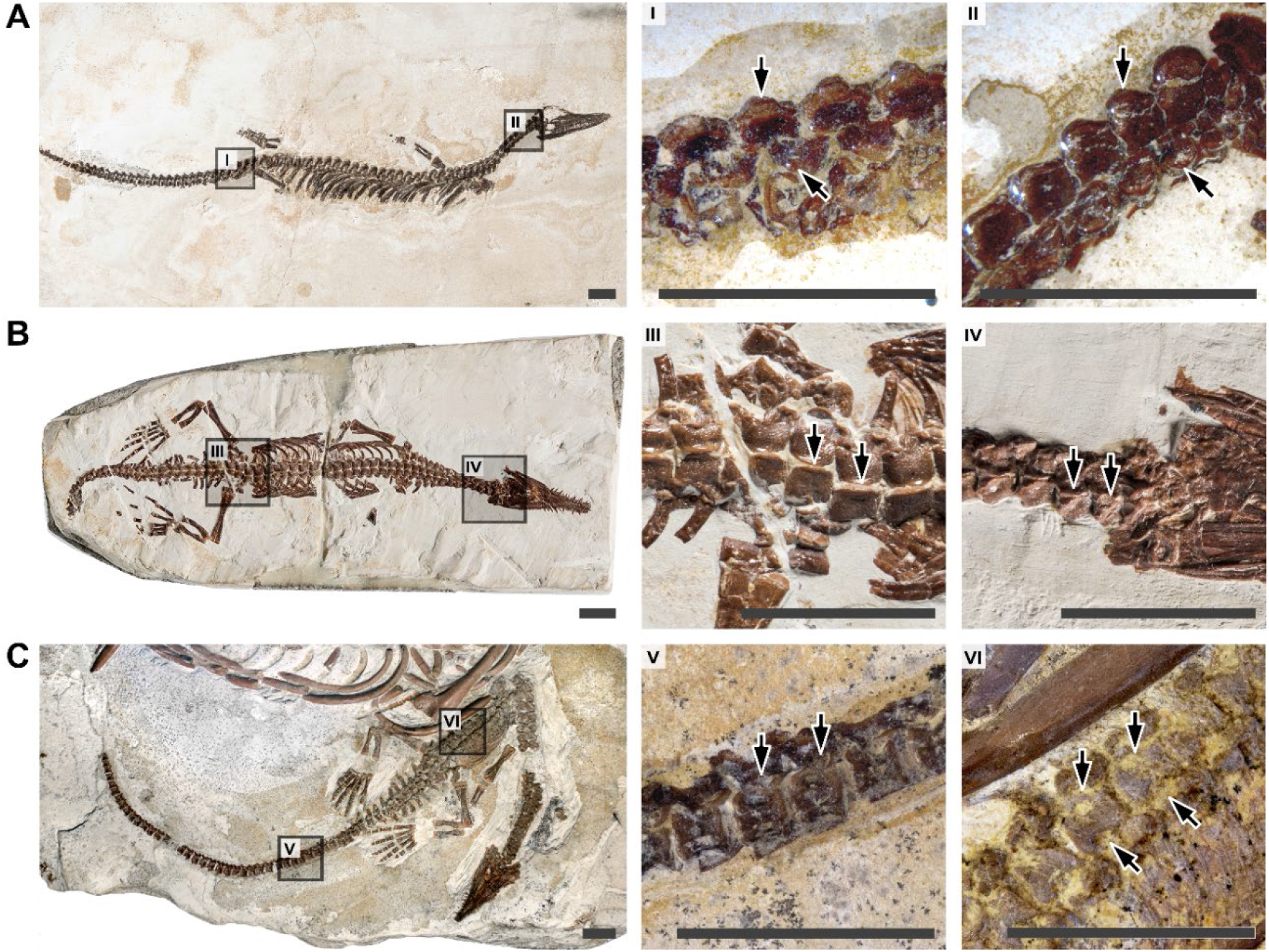
Juvenile specimens of *Stereosternum tumidum* showing axial ossification patterns. A. SMF-R-4512, showing gradients of pleurocentrum and neural arch ossification; B. BSPG 1979 I 37, showing a gradient of neural arch fusion; C. MZSP-PV 1301, showing a gradient of neurocentral fusion. Specimens are oriented with the posterior region to the left and the anterior region to the right. Dark grey squares highlight the areas shown in close-up with the matching numbering. Arrows highlight differences in degree of ossification or fusion between two adjacent close-ups. Scale bars: 10 mm.

The single anterior locus condition appears to be quite common among amniotes. Out of the 41 documented taxa, 25 present a cervical locus of ossification for the centra (Table S1). All birds and reptiles have a cervical locus, but most mammals do not. Our analysis shows that the cervical locus has been lost in mammals, with this loss happening more or less early in the history of mammals depending on the ASR model used: in Mammalia with maximum likelihood, in Theria with parsimony.

Later, a few isolated mammalian taxa *(Didelphis, Oryctolagus, Sus,* and *Tachyglossus)* reacquire the cervical locus (Table S1). With this exception, this locus is a phylogenetically constrained trait and does not seem very plastic. Thus, rather than completely disappearing, the genetic potential for a cervical locus might have been merely muted in mammals and later reactivated in the aforementioned taxa. The potential causes of this reactivation remain to be investigated, since there is no clear common denominator to these taxa.

Centrum ossification is the only axial ossification pattern we reconstruct as having two loci in amniotes ancestrally: one in the cervicals and one in the thoracics. Virtually all mammals studied have a thoracic locus of central ossification; only Talpidae do not bear the thoracic locus and have a sacral locus instead (Hautier et al., 2010). Some birds also possess an additional locus of ossification in the upper dorsal region, and this locus is reconstructed as ancestrally present in paleognathous birds (Fig. S1, S2). *Didelphis* and *Oryctolagus* present an additional lumbar locus of ossification (Table S1). We measure a very strong phylogenetic signal for each of the four loci positions (λ = 1.000, p < 0.001).

The evolutionary scenario for the thoracic locus follows a reverse path to that of the cervical locus: it is ancestrally present in amniotes, lost in reptiles but retained in mammals. The thoracic locus appears to be already absent in *Stereosternum,* which would suggest it may too have been muted or lost very early on in the history of Reptilia. After its loss, the thoracic locus is reactivated or reacquired only twice in the reptilian lineage: in paleognathous birds and in *Sterna*. While the peculiarity of development patterns in paleognaths has been pointed out (Maxwell and Larsson, 2009) and could be linked to the reacquisition of the thoracic locus, its presence in *Sterna* remains enigmatic.

The moles *Talpa* and *Mogera* are the only mammals lacking a thoracic locus and possessing a lumbar locus instead. In this case, the locus is located in the first lumbar, a little more posteriorly than in other mammals, and could be the result of a slightly shifted timing of ossification of the same overall area. However, the additional sacral locus found in the two moles is clearly distinct and unique. Prochel (2006) identified that ossification timing in Talpidae diverges significantly from the standard mammalian condition due to their fossorial lifestyle, and it is very likely that a divergence in timing would also affect axial ossification loci.

### Neural arch ossification

Much like centrum ossification, neural arch ossification in the mesosaur *Stereosternum* is progressing posteriorly from a single cervical locus (Fig. 2A). This cervical locus of neural arch ossification is present in all amniote taxa (Fig. 3B). Although it is sometimes found in association with other loci, the cervical locus is in fact the only locus that is present in all reptiles, birds, marsupials, as well as in the temnospondyl *Micromelerpeton* (Table S1).

**Figure 3.**
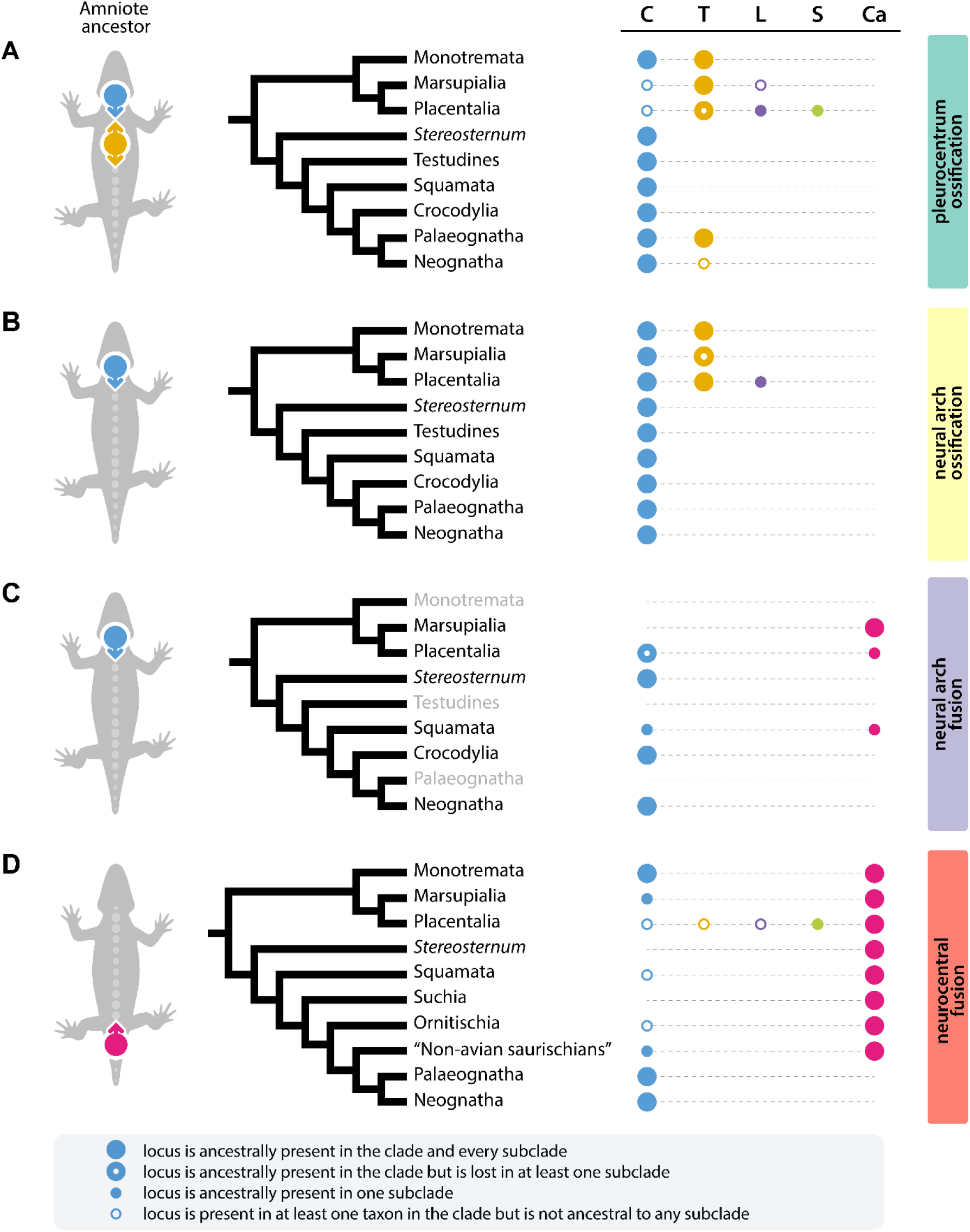
Distribution and ancestral state reconstruction of loci of axial ossification and fusion in amniotes: A. centrum ossification; B. neural arch ossification; C. neural arch fusion; D. neurocentral fusion. Abbreviations: C, cervical; T, thoracic/anterior dorsal; L, lumbar/posterior dorsal; S, sacral; Ca, caudal. Colored circles on the grey silhouette represent the reconstructed ancestral condition in amniotes and arrows show the direction of ossification/fusion from each locus. The same color code for loci positions is used all over the figure. Original data in Supplementary figures 1-4.

Additional neural arch loci are only found within Mammalia. All placentals possess a second locus of neural arch ossification in the thoracic region, as does the monotreme *Tachyglossus* (Table S1). Notably, marsupials do not have any additional neural arch loci, with the exception of *Didelphis* and *Sminthopsis* (Table S1). *Bos*, *Capra*, *Dasypus*, and *Homo* also display a third neural arch locus in the lumbar region. The distribution of the thoracic locus, which is restricted to mammals, shows a strong phylogenetic signal (λ = 0.816, p < 0.001). Similarly, we find a significant phylogenetic signal in the distribution of the lumbar locus (λ = 1.000, p = 0.004).

Our ASR recovers the single cervical locus as ancestrally present in all clades of amniotes and as the plesiomorphic condition in Amniota itself (Fig. 3B). The thoracic locus is ancestral to all clades of mammals but is lost in Australidelphia. This locus is also absent in all other non-mammalian amniotes (Fig. S3, S4). Likewise, the lumbar locus is reconstructed as ancestral in bovids but as ancestrally absent in other super-generic clades of amniotes. Parsimony mostly agrees with maximum likelihood and is only inconclusive about the ancestral presence of a thoracic locus in two clades of Australian marsupials (Fig. S4).

### Neural arch fusion

Neural arch fusion is the least well documented of the four patterns we considered. To the best of our knowledge, it has only been described in 18 extant amniotes, and never in any fossil tetrapod. Here, we document the first occurrence of this process in the mesosaur *Stereosternum.* We made our observations on three specimens of *Stereosternum:* BSPG 1979 I 37, MZSP-PV 1301, and MZSP-PV 1313. All three of them have dorsally fused neural laminae in the cervical region whereas they are still unfused in posterior parts of the body. This suggests that neural arches fuse dorsally along an anteroposterior gradient, closing like a zipper. This “zipper-like” pattern is best observed in BSPG 1979 I 37 (Fig. 2B).

Unlike centrum and neural arch ossification, all studied taxa bear only one locus of neural arch fusion, which is located in the cervical or in the caudal section of the vertebral column (Fig. 3C). Interestingly, the cervical and caudal loci seem to be mutually exclusive since we found no taxon where both conditions overlapped. Moreover, the distribution of the neural arch fusion loci among amniotes is much less phylogenetically restricted than that of the ossification loci (Fig. 3C). Among the studied reptiles, three archosaurs, two squamates, and the mesosaur *Stereosternum,* possess a cervical locus, whereas only *Bradypus*, *Homo,* and *Bos* have this locus among mammals (Table S1). The caudal locus is found in all the studied marsupials, rodents, and lagomorphs, and in some squamates (Table S1). We measure a very slight albeit non-significant phylogenetic signal for both the cervical and the caudal locus (λ = 0.471, p = 0.189). This slight phylogenetic signal is probably driven by clades displaying homogenous loci position such as Marsupialia, Glires (rodents + lagomorphs), and Archosauria (Fig. S5).

Despite homogeneity within some clades, we could not reconstruct any of the ancestral states using maximum likelihood. Only parsimony provided results (Fig. S5). The caudal locus is reconstructed as ancestral in Marsupialia and Glires (Fig. 3C), and as the most likely ancestral state in some subclades of Squamata (Fig. S5). The ancestral condition is unresolved in Squamata and its subclades. In all other clades, including Amniota, we find that the cervical locus is the probable ancestral condition (Fig. 3C).

### Neurocentral fusion

Our observations on the neurocentral fusion of *Stereosternum* are based on specimen MZSP-PV 1301. The posterior half of this individual reveals that centra and neural arches are unfused in dorsal vertebrae, whereas the posteriormost caudals are completely fused (Fig. 2C). This suggests a caudocranial (from tail to head) direction of neurocentral fusion, probably from a caudal locus.

In stark contrast to the three other ossification patterns, a cervical locus is neither the most common nor the reconstructed ancestral condition for neurocentral fusion in amniotes. Instead, neurocentral fusion ancestrally started from a caudal locus with fusion proceeding anteriorly (Fig. 3D). The caudal locus is maintained in most mammals and non-avian reptiles, but is absent in birds, rodents and lagomorphs, humans, as well as in a few squamates and dinosaurs (Table S1).

Birds deviate from the ancestral condition by adopting an anteroposterior neurocentral fusion starting from a cervical locus (Fig. 3D). The transition from a caudal to a cervical locus already occurred prior to Aves among non-avian dinosaurs: the ornithomimosaurian *Nqwebasaurus* and the tyrannosaurid *Dilong* also possess a cervical locus (Table S1), suggesting that this locus appeared in Coelurosauria (Fig. S6, S7). Moreover, both the cervical and the caudal locus are present in *Dilong*, showing that both loci coexisted in early coelurosaurians until the caudal locus later disappeared in more derived avian clades.

In addition to coelurosaurs, several mammals and dinosaurs also acquired a cervical locus in addition to the caudal one (Table S1). Moreover, certain squamates possess only the cervical locus and have lost the caudal locus. This suggests that the presence of a cervical locus is a rather plastic trait that emerged independently multiple times.

Among mammals, rodents and lagomorphs adopt a sacral locus, probably through the anterior shifting of the caudal locus (Fig. 3D). Humans are the only amniotes to present both a thoracic and a lumbar locus. It is possible that the thoracic and the lumbar locus respectively result from the posterior shifting of a cervical locus and the anterior shifting of a caudal one. Additional data on primates would be necessary for this hypothesis to be tested.

## Discussion

### Axial ossification patterns and Hox genes

Despite their evolutionary history spanning over 300 million years and their great diversity and disparity, amniotes are surprisingly conservative with respect to axial ossification patterns. This is also reflected in the significant phylogenetic signal recovered in each of the patterns (Fig. S1-S7). It is remarkable that in the vast majority of amniotes some loci as well as the directionality of axial ossification has been maintained since the initial evolution of Amniota (Fig. 3). This is particularly interesting considering the disparity of vertebral morphologies and functions in the clade, and the diverse patterns and modes of axial regionalization that have been recognized (Böhmer et al., 2018; Bui and Larsson, 2021; Jones et al., 2018; Krings et al., 2014; Terray et al., 2020).

It seems likely that the observed conservatism of axial ossification patterns is due to constraints imposed by the underlying genetic processes during vertebral development. While genetic material does not preserve in deep geological time, comparisons with modern representatives can provide some insights into the genetic underpinnings of axial patterning in fossil taxa (Böhmer et al., 2015; Head and Polly, 2015; Müller et al., 2010). In this context, the role of *Hox* genes and their respective regulators in directing axial patterning in vertebrates specifically and animals more broadly, has long been known (Böhmer et al., 2015; Head and Polly, 2007; McGinnis and Krumlauf, 1992; Shubin et al., 2009; Wellik, 2007).

Moreover, a number of studies have shown that overlapping expression domains of *Hox* genes along the anteroposterior axis of the embryo directly impact vertebral morphologies in the adult (Böhmer, 2017; Böhmer et al., 2015; Burke et al., 1995; Galis, 1999; Hautier et al., 2014, 2010; Ohya et al., 2005) and that the ranges of *Hox* gene expression domains can also influence the relative timing of ossification in vertebral elements (Bui and Larsson, 2021; Hautier et al., 2014, 2010). Therein, certain *Hox4-10* paralogues linked to vertebral regionalization have been suggested to have been retained in Amniota since the last common ancestor of the clade (Head and Polly, 2015). Therefore, axial ossification models including those of fossil taxa may indeed reflect to some degree how developmental constraints prevented or channelized evolutionary innovation. However, the more detailed roles that *Hox* and other genes play in directing ossification and fusion patterns in extinct and extant taxa alike remains to be investigated.

Among the main clades of amniotes, two stand out in terms of these patterns: mammals notably diverge from the ancestral amniote condition for centrum and neural arch ossification, and to a lesser extent, birds also diverge from the ancestral amniotic neurocentral fusion pattern. In addition, several subclades within mammals and birds diverge from the ancestral condition of their parent taxa: Paleognathae and Talpidae for centrum ossification, Australidelphia for neural arch ossification, Marsupialia for neural arch fusion, Glires for neural arch fusion and neurocentral fusion (Fig. 3). This suggests that mammals and birds have a greater potential for variation in their axial ossification patterns than the other main groups of amniotes, like squamates and turtles.

Mammals and birds have a strongly regionalized vertebral column, which means that their column is segmented into more morphologically and functionally distinct regions than other amniotes (Böhmer et al., 2018; Bui and Larsson, 2021; Jones et al., 2020, 2018; Krings et al., 2014; Terray et al., 2020). Strongly regionalized columns result from complex interactions between *Hox* genes (Burke et al., 1995; Young et al., 2009), and changes in *Hox* gene expression domains can in turn influence the timing of ossification (Bui and Larsson, 2021; Hautier et al., 2014, 2010). With their strongly regionalized columns, mammals and birds possess a more strongly modularized vertebral column, which in turn may provide a greater potential for the accumulation of changes within each module and in the overall patterning and timing of ossification (Jones et al., 2019). Therefore, the greater diversity of axial ossification patterns in mammals and birds may directly results from their regionalization.

Although the current state of knowledge prevents us from statistically testing the links between regionalization and axial ossification, there are a few cases where changes in regionalization parallel modifications of axial ossification patterns. For instance, the transition in coelurosaurian dinosaurs from a caudal to a cervical locus of neurocentral fusion strikingly mirrors the progressive loss of articulation in tail vertebrae in favor of a fused pygostyle in birds, a loss probably caused itself by variations in *Hox* genes (Rashid et al., 2014). Similarly, the adoption of a fixed number of seven cervicals in mammals through *Hox* gene mutations coincides with the acquisition of an additional thoracic locus of neural arch ossification (Böhmer, 2017; Müller et al., 2010).

While these examples are consistent with our hypothesis and the study of vertebral regionalization has expanded rapidly in recent years (Bui and Larsson, 2021; Jones et al., 2020, 2019; Terray et al., 2020), some amniote clades remain under-represented in terms of quantitative data on the topic. For instance, our hypothesis predicts that non-avian reptiles should have less complex patterns of *Hox* gene expression during axial development matching the poorly regionalized vertebral column in the adults. Determining whether this is indeed the case would provide more insights into potential links between these two processes. This shows how investigating vertebral regionalization and development in further non-model amniote taxa would be crucial to understand the evolutionary history of the vertebral column in amniotes.

### Scenarios for the evolution of axial ossification patterns

We propose the following ancestral condition for axial ossification patterns in amniotes: (i) pleurocentrum ossification proceeding posteriorly from two loci in the cervical and the thoracic region, (ii) neural arch ossification proceeding posteriorly from a single cervical locus, (iii) neural arch fusion progressing posteriorly from a single cervical locus in a “zipper-like” pattern, and (iv) neurocentral fusion proceeding anteriorly from a caudal locus. From this ancestral condition, their exact evolutionary development remains obscure due to the current lack of fossil data. On the reptilian branch of Amniota, the exceptional specimens of *Stereosternum* provide unique data from the fossil record that fill the gap that data based on extant animals alone leave behind. For each axial ossification pattern, *Stereosternum* displays the reconstructed ancestral reptilian condition, and for all patterns but centrum ossification, it even displays the ancestral amniote condition. The observations in this ancient amniote taxon therefore lend some support to our reconstruction.

On the mammalian branch however, the lack of information for vertebral ossification patterns in non-mammalian synapsids is problematic. Axial ossification patterns have never been documented in any pelycosaur-grade synapsid in therapsids, in Mesozoic or even Cenozoic mammals. Additional data from fossil synapsids would greatly help to refine the resolution and robustness of the reconstructed evolutionary history of these traits in amniotes.

Based on the current data and depending on what will eventually be found in synapsids in the future, we predict three scenarios: (i) fossil synapsids show patterns of axial ossification resembling those of reptiles. In that case, the observed reptilian patterns would reflect the ancestral condition in all amniotes. The condition observed in modern mammals would then constitute a synapomorphy adopted relatively late in their evolutionary history. This would support our speculation that the specific axial ossification patterns found in crown-mammals may be connected to the evolution of a stronger vertebral regionalization in therapsids (see also Jones et al., 2020); (ii) fossil synapsids show patterns of axial ossification resembling those of crown-mammals. This would imply that the two main branches of Amniota each developed their own specific modes of axial ossification very early in their evolutionary history, potentially at the time of their original dichotomy. This would in return suggest that axial ossification patterns might have played a role in this dichotomy; (iii) fossil synapsids show unique patterns of axial ossification. This scenario would combine the other two, with a dichotomy in axial ossification patterns at the base of Amniota and the innovation of another, different condition in crown-mammals. Regardless of which scenario might be supported once additional data on fossil synapsids becomes available, it will yield important implications for the evolutionary history of morphological diversity in amniotes.

## Conclusions

Reviewing the literature and with additional data from exceptionally well-preserved fossils, we reconstruct the ancestral axial ossification and fusion patterns in amniotes:

- Centra ossify from neck to tail, starting from two loci in the cervical and the thoracic region.
- Neural arches also ossify posteriorly but from a single cervical locus.
- Neural arches fuse together in a “zipper-like” pattern, starting in a cervical locus and progressing posteriorly.
- Neurocentral fusion begins in the caudal region and progresses anteriorly towards the head.

Despite the long evolutionary history of amniotes, all four axial ossification patterns show a strong phylogenetic signal and appear to have been quite stable over time. We propose that this conservatism may be linked to constraints posed by underlying developmental processes across amniotes. Our study also highlights variability in the axial ossification patterns of birds and mammals. We suggest a correlation between this variability and the strong vertebral regionalization existing in these clades. This study provides a framework for further research on axial ossification in fossil and modern amniotes, but more fossil and molecular data will be necessary to further resolve the ancestral ossification patterns within Amniota.

## Materials and Methods

Our study focuses on four major patterns of axial ossification: centrum ossification, neural arch ossification, neural arch fusion, and neurocentral fusion. For each of these patterns, we reviewed the existing literature on extant and fossil amniotes and gathered information on two key aspects: the number of loci in the spine from where the patterns start and the position of these loci in the vertebral column. Very little information was available on these patterns in the fossil record. The axial skeleton of juvenile specimens of extinct amniotes has rarely been described in sufficient detail for the scope of our study.

Fortunately, based on exceptional specimens of the early Permian mesosaurid reptile *Stereosternum tumidum,* we were able document patterns of axial ossification in one of the most basal clades of amniotes. In addition, to complete missing information on the patterns of some extant taxa, observations made on specimens from the collections of the Museum für Naturkunde in Berlin, Germany were included (Table S1).

A composite tree of all studied species was constructed in Mesquite 3.6(Maddison and Maddison, 2019) combining recently published phylogenies (Fig. S8). In addition to the taxa for which vertebral ossification data was available, fossil taxa were included in the tree to obtain a more accurate time-calibration following recommendations from the Fossil Calibration Database (Ksepka et al., 2015). The resulting tree was time-calibrated based on occurrence dates from the Paleobiology Database (Uhen and Sessa, 2013) and using equal time-scaling method. A subset of the time-calibrated tree was then generated for each of the four ossification patterns by trimming the original tree to match the taxon sample with available data.

In each studied taxon, the four ossification patterns were characterized in terms of number of loci and the position of these loci in the vertebral column. The presence or absence of a locus was noted in five sections of the vertebral column for each pattern: (i) cervical, (ii) upper dorsal/thoracic, (iii) lower dorsal/lumbar (iv) sacral, and (v) caudal. The presence/absence of a locus in these sections was scored as a binary character. As a result, each ossification pattern was decomposed into a matrix of four binary characters, one for each section of the vertebral column (Table S1). The binary characters were then tested for phylogenetic signal using Pagel’s λ (Pagel, 1999). Ancestral state reconstruction was performed on each character twice, once using maximum likelihood with symmetrical rates of evolution, and once using maximum parsimony with sequential transition costs.

Tree calibration and trimming, statistical analyses, and ancestral character reconstructions were performed using the ape (Paradis and Schliep, 2019), castor (Louca and Doebeli, 2017), paleotree (Bapst, 2012), and phytools (Revell, 2012) packages in R v.4.0.5 (R. Core Team, 2021). Code in supplements (Dataset S1).

## Acknowledgments

We thank R. Brocke (Senckenberg Naturmuseum, Frankfurt), A. Carvalho (Museu de Zoologia da Universidade de São Paulo, SP), P. Eckhoff (Museum für Naturkunde, Berlin), C. Funk (Museum für Naturkunde, Berlin), O. Rauhut (Bayerische Staatssammlung für Paläontologie und Geologie, Munich), M.-O. Rödel (Museum für Naturkunde, Berlin) for access to collections and specimen loans. Very special thanks to A. Carvalho and H. Zaher (MZSP) who helped us obtain crucial photographs of MZSP despite the pandemic. Additional thanks to J. Müller, M.J MacDougall and Y. Haridy (Museum für Naturkunde, Berlin) for discussion and their helpful comments during the drafting of this article. This work was supported by the German Research Foundation (DFG FR 2457/6-1).

