## Supplementary Information for "Regionalization, constraints, and the ancestral ossification patterns in the vertebral column of amniotes"

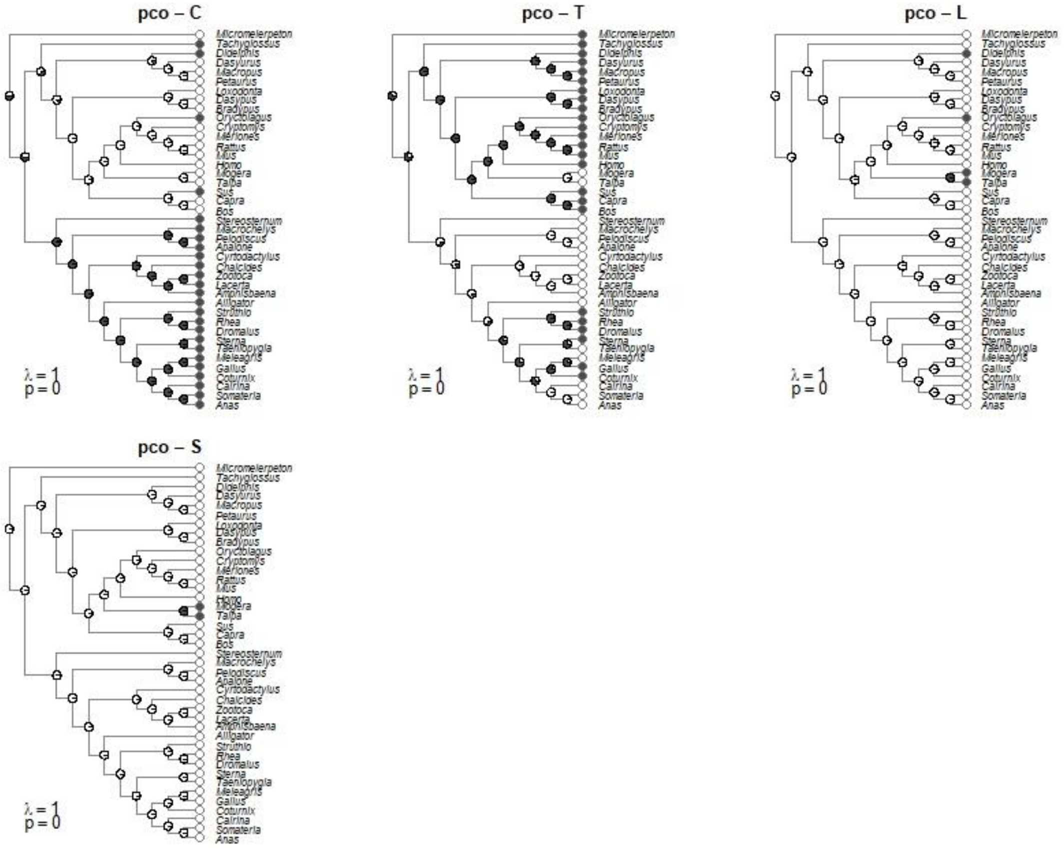

**Fig. S1.** Ancestral state reconstruction of the presence of a locus for pleurocentrum ossification in each section of the vertebral column, using maximum likelihood. In Fig. S1 to S8, sections are labelled as follows: C: cervical; T: thoracic/upper dorsal; L: lumbar/lower dorsal; S: sacral; Ca: caudal. The four patterns are abbreviated as follows: PCO: pleurocentrum ossification; NAO: neural arch ossification, NAF: neural arch fusion, NCF: neurocentral fusion. Black and white circles at tips respectively mark the presence and the absence of a locus in the section. Pie charts at nodes show the probability of each state being ancestral for the clade.

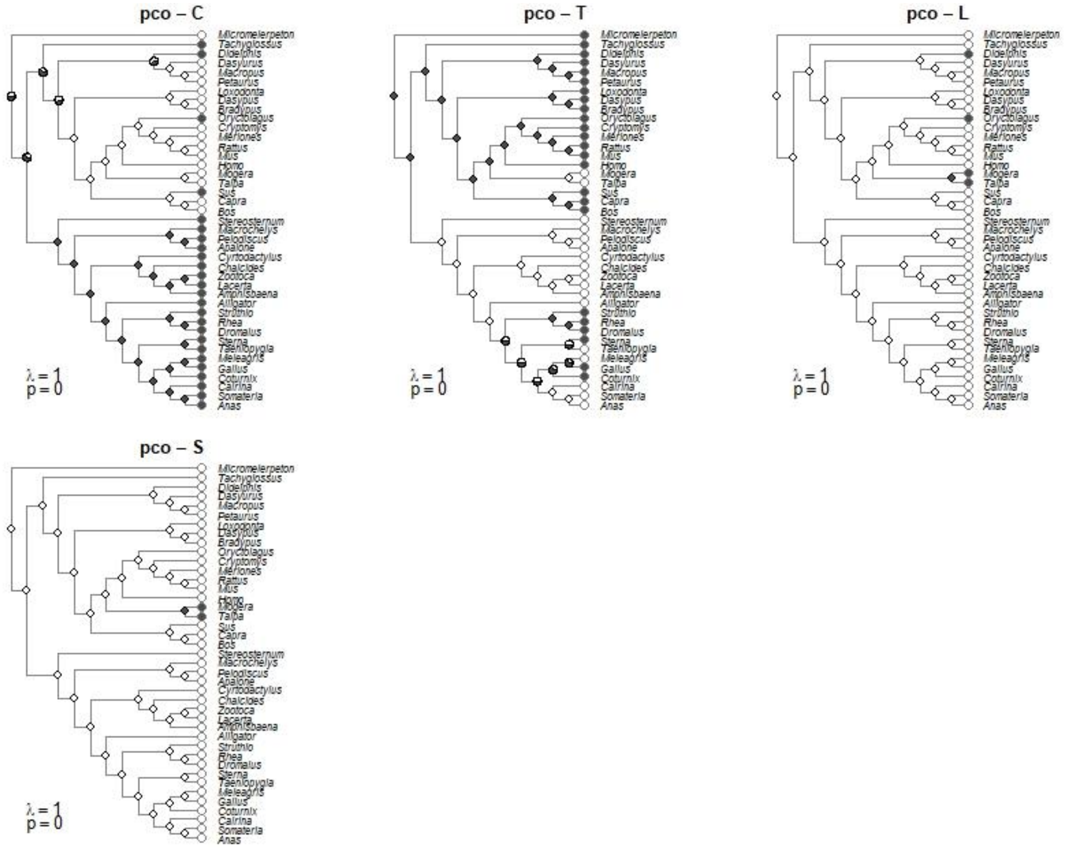

**Fig. S2.** Ancestral state reconstruction of the presence of a locus for pleurocentrum ossification in each section of the vertebral column, using parsimony.

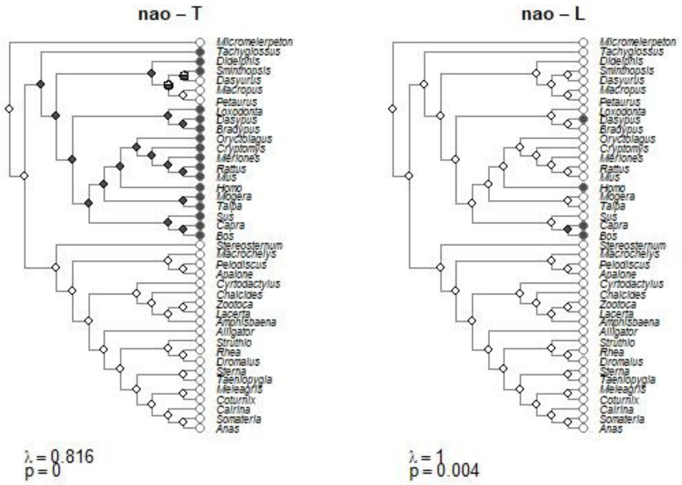

**Fig. S3.** Ancestral state reconstruction of the presence of a locus for neural arch ossification in each section of the vertebral column, using maximum likelihood.

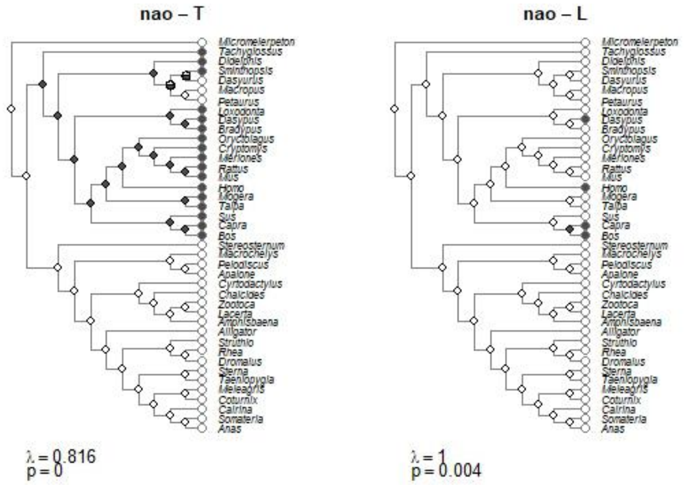

**Fig. S4.** Ancestral state reconstruction of the presence of a locus for neural arch ossification in each section of the vertebral column, using parsimony.

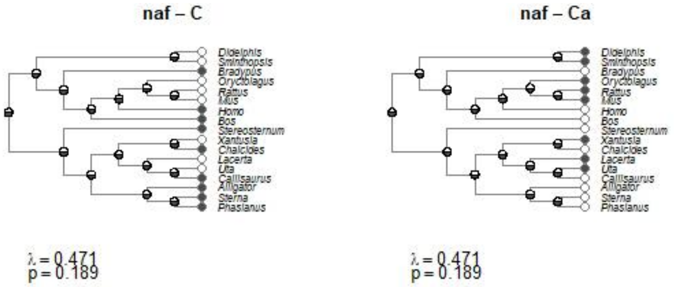

**Fig. S5.** Ancestral state reconstruction of the presence of a locus for neural arch fusion in each section of the vertebral column, using maximum likelihood.

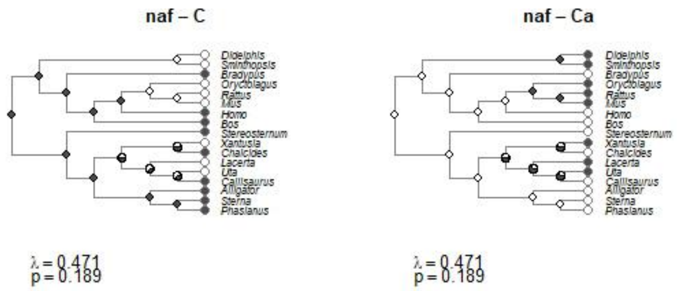

**Fig. S6.** Ancestral state reconstruction of the presence of a locus for neural arch fusion in each section of the vertebral column, using parsimony.

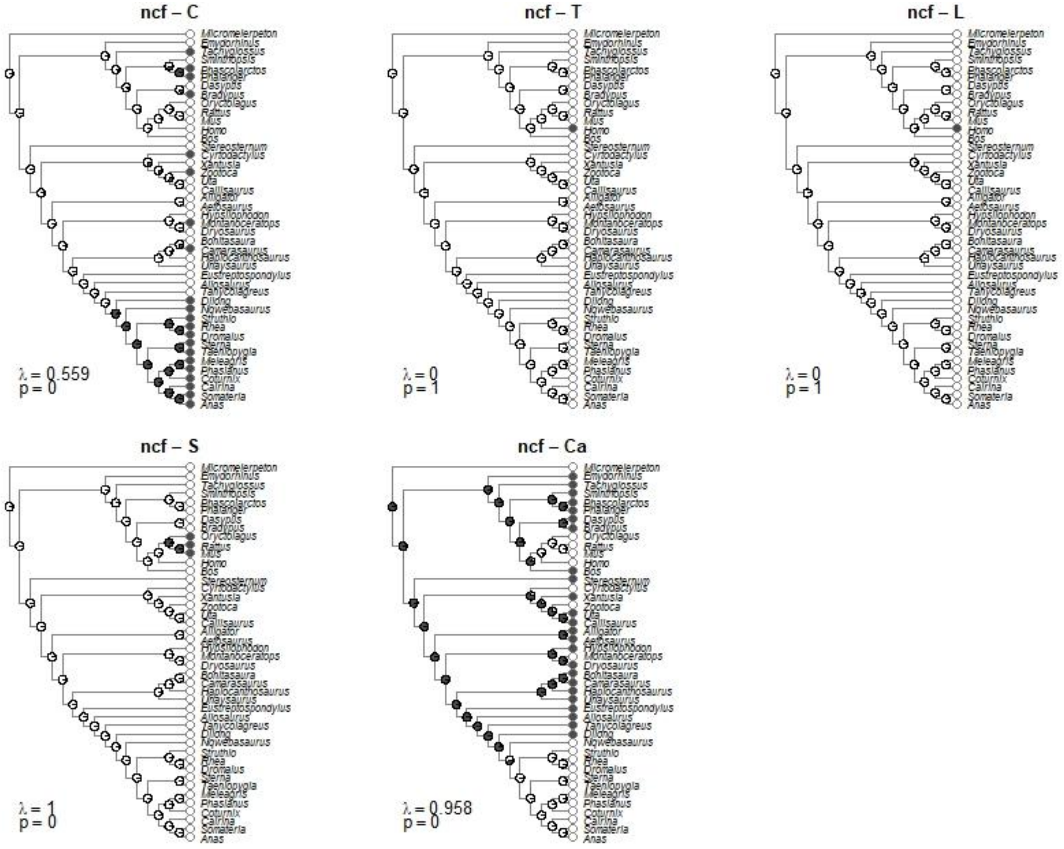

**Fig. S7.** Ancestral state reconstruction of the presence of a locus for neurocentral fusion in each section of the vertebral column, using maximum likelihood.

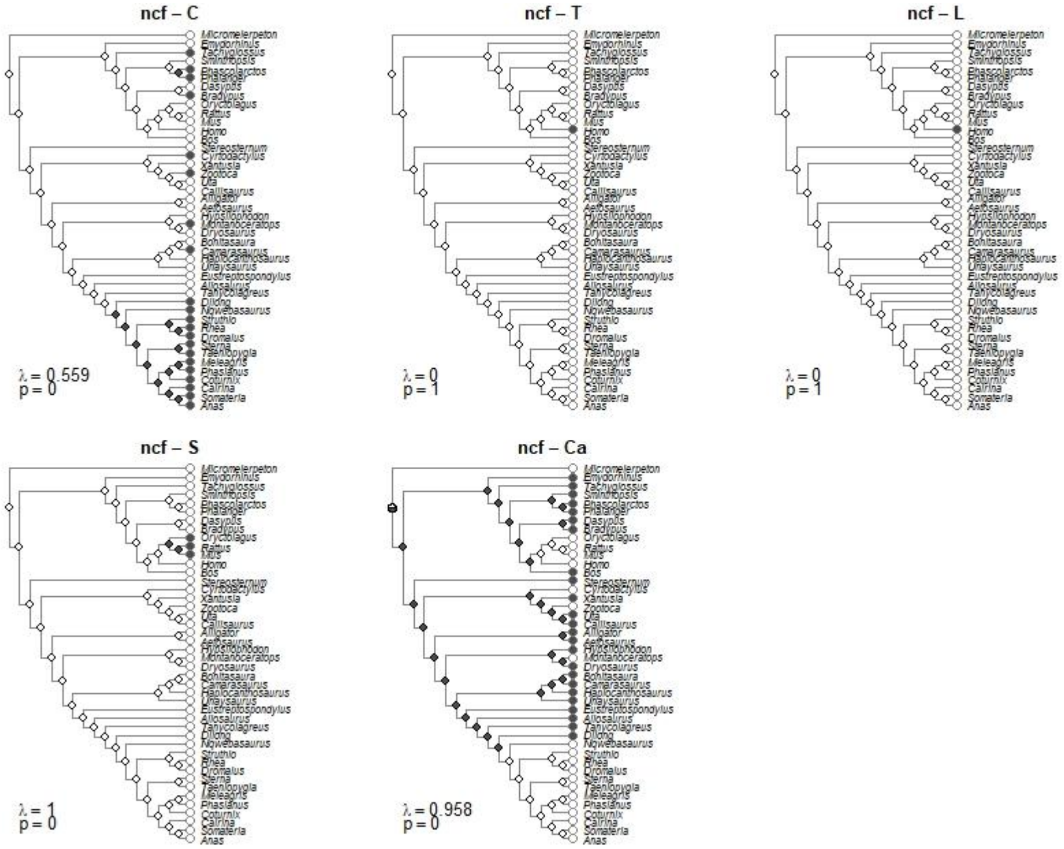

**Fig. S8.** Ancestral state reconstruction of the presence of a locus for neurocentral fusion in each section of the vertebral column, using parsimony.

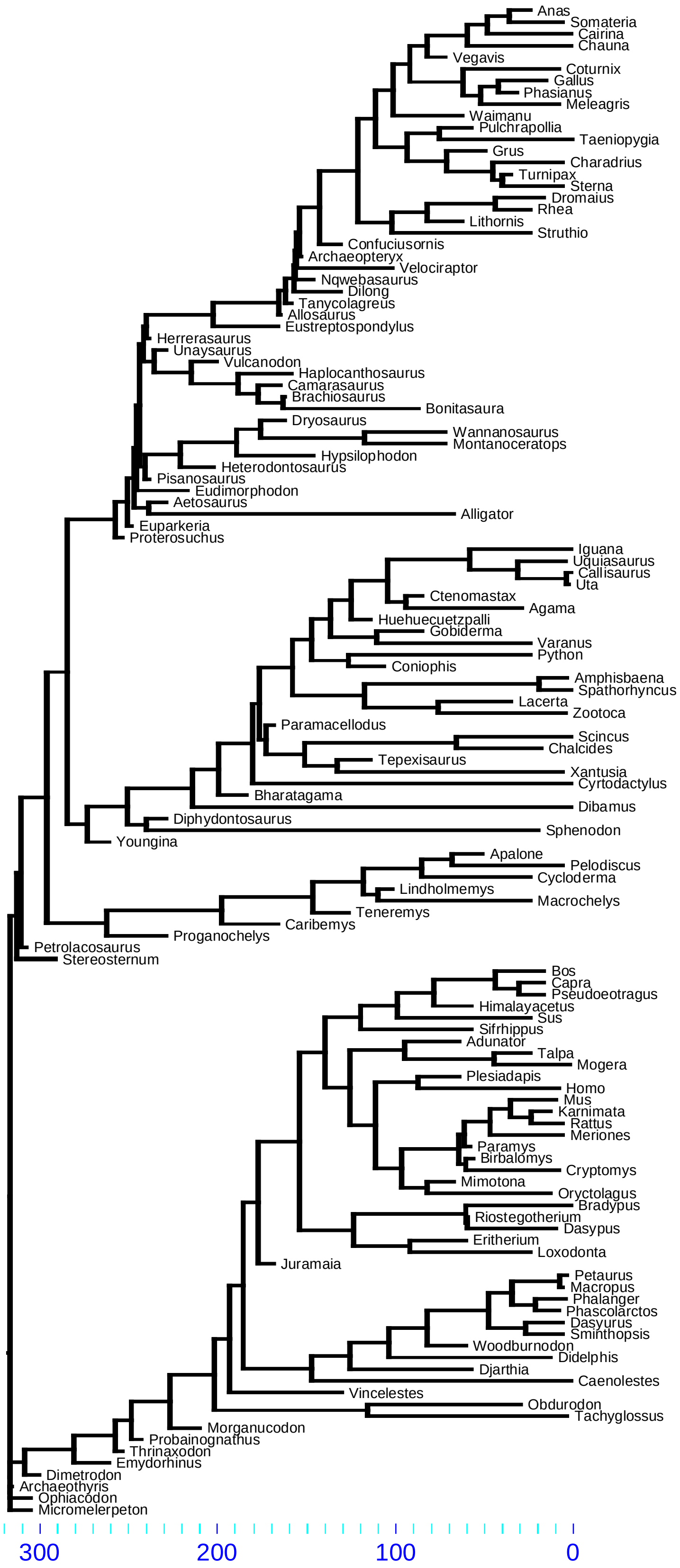

**Fig. S9.** Time-calibrated tree of amniotes using combined topographies **(1–14)**. Ages in Ma.

**Table S1.** Position of ossification and fusion loci in the studied taxa. Sections of the vertebral column are labelled as follows: C: cervical; T: thoracic/upper dorsal; L: lumbar/lower dorsal; S: sacral; Ca: caudal. Black and white circles respectively mark the presence and absence of a locus in the vertebral region. Hyphens marks the absence of data for this pattern.

| Taxa | References / specimens | PCO | | | | | NAO | | | | | NAF | | | | | NCF | | | | |
|---|---|---|---|---|---|---|---|---|---|---|---|---|---|---|---|---|---|---|---|---|---|
| | | C | T | L | S | Ca | C | T | L | S | Ca | C | T | L | S | Ca | C | T | L | S | Ca |
| *†Aetosaurus* | (15) | - | - | - | - | - | - | - | - | - | - | - | - | - | - | - | ○ | ○ | ○ | ○ | ● |
| *Alligator* | (16–18) | ● | ○ | ○ | ○ | ○ | ● | ○ | ○ | ○ | ○ | ● | ○ | ○ | ○ | ○ | ○ | ○ | ○ | ○ | ● |
| *†Allosaurus* | (15, 19) | - | - | - | - | - | - | - | - | - | - | - | - | - | - | - | ○ | ○ | ○ | ○ | ● |
| *Amphisbaena* | (20) | ● | ○ | ○ | ○ | ○ | ● | ○ | ○ | ○ | ○ | - | - | - | - | - | - | - | - | - | - |
| *Anas* | (21, 22) | ● | ○ | ○ | ○ | ○ | ● | ○ | ○ | ○ | ○ | - | - | - | - | - | ● | ○ | ○ | ○ | ○ |
| *Apalone* | (23) | ● | ○ | ○ | ○ | ○ | ● | ○ | ○ | ○ | ○ | - | - | - | - | - | - | - | - | - | - |
| *†Bonitasaura* | (24) | - | - | - | - | - | - | - | - | - | - | - | - | - | - | - | ○ | ○ | ○ | ○ | ● |
| *Bos* | (25) ZMB.Mam.108963 | ○ | ● | ○ | ○ | ○ | ● | ● | ● | ○ | ○ | ● | ○ | ○ | ○ | ○ | ○ | ○ | ○ | ○ | ● |
| *Bradypus* | (26) ZMB.Mam.106949 ZMB.Mam.102630 | ○ | ● | ○ | ○ | ○ | ● | ● | ○ | ○ | ○ | ● | ○ | ○ | ○ | ○ | ● | ○ | ○ | ○ | ● |
| *Cairina* | (21) | ● | ○ | ○ | ○ | ○ | ● | ○ | ○ | ○ | ○ | - | - | - | - | - | ● | ○ | ○ | ○ | ○ |
| *Callisaurus* | (27) | - | - | - | - | - | - | - | - | - | - | ● | ○ | ○ | ○ | ○ | ○ | ○ | ○ | ○ | ● |
| *†Camarasaurus* | (28) | - | - | - | - | - | - | - | - | - | - | - | - | - | - | - | ● | ○ | ○ | ○ | ● |
| *Capra* | (26) | ○ | ● | ○ | ○ | ○ | ● | ● | ● | ○ | ○ | - | - | - | - | - | - | - | - | - | - |
| *Chalcides* | (29) | ● | ○ | ○ | ○ | ○ | ● | ○ | ○ | ○ | ○ | ● | ○ | ○ | ○ | ○ | - | - | - | - | - |
| *Coturnix* | (22, 30) | ● | ● | ○ | ○ | ○ | ● | ○ | ○ | ○ | ○ | - | - | - | - | - | ● | ○ | ○ | ○ | ○ |
| *Cryptomys* | (26) | ○ | ● | ○ | ○ | ○ | ● | ● | ○ | ○ | ○ | - | - | - | - | - | - | - | - | - | - |
| *Cyrtodactylus* | (31) | ● | ○ | ○ | ○ | ○ | ● | ○ | ○ | ○ | ○ | - | - | - | - | - | ● | ○ | ○ | ○ | ○ |
| *Dasypus* | (26) ZMB.Mam.85925 | ○ | ● | ○ | ○ | ○ | ● | ● | ● | ○ | ○ | - | - | - | - | - | ○ | ○ | ○ | ○ | ● |
| *Dasyurus* | (26) | ○ | ● | ○ | ○ | ○ | ● | ○ | ○ | ○ | ○ | - | - | - | - | - | - | - | - | - | - |
| *Didelphis* | (32, 33) | ● | ● | ● | ○ | ○ | ● | ● | ○ | ○ | ○ | ○ | ○ | ○ | ○ | ● | - | - | - | - | - |
| *†Dilong* | (34) | - | - | - | - | - | - | - | - | - | - | - | - | - | - | - | ● | ○ | ○ | ○ | ● |
| *Dromaius* | (35) | ● | ● | ○ | ○ | ○ | ● | ○ | ○ | ○ | ○ | - | - | - | - | - | ● | ○ | ○ | ○ | ○ |
| *†Dryosaurus* | (36) | - | - | - | - | - | - | - | - | - | - | - | - | - | - | - | ○ | ○ | ○ | ○ | ● |
| *Emydorhinus* | (37) | - | - | - | - | - | - | - | - | - | - | - | - | - | - | - | ○ | ○ | ○ | ○ | ● |
| *†Eustreptospondylus* | (15, 38) | - | - | - | - | - | - | - | - | - | - | - | - | - | - | - | ○ | ○ | ○ | ○ | ● |
| *Gallus* | (39) | ● | ● | ○ | ○ | ○ | - | - | - | - | - | - | - | - | - | - | - | - | - | - | - |
| *†Haplocanthosaurus* | (15) | - | - | - | - | - | - | - | - | - | - | - | - | - | - | - | ○ | ○ | ○ | ○ | ● |
| *Homo* | (40–45) | ○ | ● | ○ | ○ | ○ | ● | ● | ● | ○ | ○ | ● | ○ | ○ | ○ | ○ | ○ | ● | ● | ○ | ○ |
| *†Hypsilophodon* | (36) | - | - | - | - | - | - | - | - | - | - | - | - | - | - | - | ○ | ○ | ○ | ○ | ● |
| *Lacerta* | (46) | ● | ○ | ○ | ○ | ○ | ● | ○ | ○ | ○ | ○ | ○ | ○ | ○ | ○ | ● | - | - | - | - | - |

| | | | | | | | | | | | | | | | | | | | | | |
|---|---|---|---|---|---|---|---|---|---|---|---|---|---|---|---|---|---|---|---|---|---|
| *Loxodonta* | (26) | ○ | ● | ○ | ○ | ○ | ● | ● | ○ | ○ | ○ | - | - | - | - | - | - | - | - | - | - |
| *Macrochelys* | (47) | ● | ○ | ○ | ○ | ○ | ● | ○ | ○ | ○ | ○ | - | - | - | - | - | - | - | - | - | - |
| *Macropus* | (26) | ○ | ● | ○ | ○ | ○ | ● | ○ | ○ | ○ | ○ | - | - | - | - | - | - | - | - | - | - |
| *Meleagris* | (48) | ● | ○ | ○ | ○ | ○ | ● | ○ | ○ | ○ | ○ | - | - | - | - | - | ● | ○ | ○ | ○ | ○ |
| *Meriones* | (49) | ○ | ● | ○ | ○ | ○ | ● | ● | ○ | ○ | ○ | - | - | - | - | - | - | - | - | - | - |
| †*Micromelerpeton* | (50) | ○ | ● | ○ | ○ | ○ | ● | ○ | ○ | ○ | ○ | - | - | - | - | - | ○ | ○ | ○ | ○ | ○ |
| *Mogera* | (26) | ○ | ○ | ● | ● | ○ | ● | ● | ○ | ○ | ○ | - | - | - | - | - | - | - | - | - | - |
| †*Montanoceratops* | (51) | - | - | - | - | - | - | - | - | - | - | - | - | - | - | - | ● | ○ | ○ | ○ | ○ |
| *Mus* | (52, 53) | ○ | ● | ○ | ○ | ○ | ● | ● | ○ | ○ | ○ | ○ | ○ | ○ | ○ | ● | ○ | ○ | ○ | ● | ○ |
| †*Nqwebasaurus* | (15, 54) | - | - | - | - | - | - | - | - | - | - | - | - | - | - | - | ● | ○ | ○ | ○ | ○ |
| *Oryctolagus* | (55)<br>ZMB.Mam.An.18299 | ● | ● | ● | ○ | ○ | ● | ● | ○ | ○ | ○ | ○ | ○ | ○ | ○ | ● | ○ | ○ | ○ | ● | ○ |
| *Pelodiscus* | (56) | ● | ○ | ○ | ○ | ○ | ● | ○ | ○ | ○ | ○ | - | - | - | - | - | - | - | - | - | - |
| *Petaurus* | (26) | ○ | ● | ○ | ○ | ○ | ● | ○ | ○ | ○ | ○ | - | - | - | - | - | - | - | - | - | - |
| *Phalanger* | ZMB.Mam.35319 | - | - | - | - | - | - | - | - | - | - | - | - | - | - | - | ● | ○ | ○ | ○ | ● |
| *Phascolarctos* | ZMB.Mam.36038 | - | - | - | - | - | - | - | - | - | - | - | - | - | - | - | ● | ○ | ○ | ○ | ● |
| *Phasianus* | Pers. obs. | - | - | - | - | - | - | - | - | - | - | ● | ○ | ○ | ○ | ○ | ● | ○ | ○ | ○ | ○ |
| *Rattus* | (57)<br>ZMB.Mam.11416 | ○ | ● | ○ | ○ | ○ | ● | ● | ○ | ○ | ○ | ○ | ○ | ○ | ○ | ● | ○ | ○ | ○ | ● | ○ |
| *Rhea* | (35) | ● | ● | ○ | ○ | ○ | ● | ○ | ○ | ○ | ○ | - | - | - | - | - | ● | ○ | ○ | ○ | ○ |
| *Sminthopsis* | (58) | - | - | - | - | - | ● | ● | ○ | ○ | ○ | ○ | ○ | ○ | ○ | ● | ○ | ○ | ○ | ○ | ● |
| *Somateria* | (21) | ● | ○ | ○ | ○ | ○ | ● | ○ | ○ | ○ | ○ | - | - | - | - | - | ● | ○ | ○ | ○ | ○ |
| †*Stereosternum* | BSPG 1979 I 37<br>MZSP-PV 1301<br>MZSP-PV 1313<br>SMF-R-4512 | ● | ○ | ○ | ○ | ○ | ● | ○ | ○ | ○ | ○ | ● | ○ | ○ | ○ | ○ | ○ | ○ | ○ | ○ | ● |
| *Sterna* | (59) | ● | ● | ○ | ○ | ○ | ● | ○ | ○ | ○ | ○ | ● | ○ | ○ | ○ | ○ | ● | ○ | ○ | ○ | ○ |
| *Struthio* | (35) | ● | ● | ○ | ○ | ○ | ● | ○ | ○ | ○ | ○ | - | - | - | - | - | ● | ○ | ○ | ○ | ○ |
| *Sus* | (60) | ● | ● | ○ | ○ | ○ | ● | ● | ○ | ○ | ○ | - | - | - | - | - | - | - | - | - | - |
| *Tachyglossus* | (61)<br>ZMB.Mam.35994 | ● | ● | ○ | ○ | ○ | ● | ● | ○ | ○ | ○ | - | - | - | - | - | ● | ○ | ○ | ○ | ● |
| *Taeniopygia* | (22) | ● | ○ | ○ | ○ | ○ | ● | ○ | ○ | ○ | ○ | - | - | - | - | - | ● | ○ | ○ | ○ | ○ |
| *Talpa* | (26) | ○ | ○ | ● | ● | ○ | ● | ● | ○ | ○ | ○ | - | - | - | - | - | - | - | - | - | - |
| †*Tanycolagreus* | (15, 62) | - | - | - | - | - | - | - | - | - | - | - | - | - | - | - | ○ | ○ | ○ | ○ | ● |
| †*Unaysaurus* | (15) | - | - | - | - | - | - | - | - | - | - | - | - | - | - | - | ○ | ○ | ○ | ○ | ● |
| *Uta* | (27) | - | - | - | - | - | - | - | - | - | - | ○ | ○ | ○ | ○ | ● | ○ | ○ | ○ | ○ | ● |
| *Xantusia* | (63) | - | - | - | - | - | - | - | - | - | - | ○ | ○ | ○ | ○ | ● | ○ | ○ | ○ | ○ | ● |
| *Zootoca* | (64) | ● | ○ | ○ | ○ | ○ | ● | ○ | ○ | ○ | ○ | - | - | - | - | - | ● | ○ | ○ | ○ | ○ |

**Dataset S1 (separate file).** R code and files used for the ancestral state reconstruction and the statistical analysis.
