## Supplementary Dataset 1 for "Regionalization, constraints, and the ancestral ossification patterns in the vertebral column of amniotes": ASR_Rcode.R

```
library(ape)
library(paleotree)
library(phytools)
library(reshape2)
library(Rmisc)

###>>>  Functions  <<<###

"asr" <- function(x, taxa, phy, model = c("ER", "SYM")){
  ## this function applies a maximum likelihood ASR to a trimmed tree
  require(phytools)
  # trimming NA
  d <- x
  names(d) <- taxa
  d <- d[!is.na(d)]
  tree <- drop.tip(phy, taxa[is.na(x)])
  # asr
  fit <- rerootingMethod(tree, d, model = model)
  res <- list(tree = tree, asr = fit, data = d)
  class(res) <- "asr"
  return(res)
}

"mpr" <- function(x, taxa, phy, mod){
  ## this function applies a parsimony ASR to a trimmed tree
  require(ape)
  require(castor)
  # trimming NA
  d <- x
  names(d) <- taxa
  d <- d[!is.na(d)]
  d2 <- as.integer(as.character(d))
  vmin <- min(d2)
  vmax <- max(d2)
  if(min(d2, na.rm = T) == 0) d2 <- d2 + 1
  tree <- drop.tip(phy, taxa[is.na(x)])
  # asr
  fit <- asr_max_parsimony(tree, d2, transition_costs = mod)
  lik <- as.data.frame(fit$ancestral_likelihoods)
  colnames(lik) <- vmin:vmax
  lik <- lik[sapply(lik, sum) != 0]
  fit$lik <- as.matrix(lik)
  class(fit) <- "mpr"
  res <- list(tree = tree, asr = fit, data = d)
  class(res) <- "asr"
  return(res)
}

"phylosig.asr" <- function(x){
  require(phytools)
  ## this function measures the strength of the phylogenetic signal in asr objects
  if(!inherits(x, "asr"))
    stop("x must be an object of class 'asr'")
  char <- as.numeric(x$data)
  names(char) <- names(x$data)
  L <- phylosig(x$tree, char, test = T, method = "lambda", nsim = 10000)$lambda
  p <- phylosig(x$tree, char, test = T, method = "lambda", nsim = 10000)$P
  res <- c(lambda = L, p = p)
  return(res)
}

"plot.asr" <- function(x, n, phy = x$tree, signal = T, 
                       tip.size = 1.5, tip.col = "gray50",  pie.size = .6, pie.col = "black", y.lim = 40, 
                       palette = c("#4d4d4d", "#ffffff", "#189e77", "#4e9bd4", "#e62a8a", "#7866ac", "#a6d056", "#a6761d","#e5ac23"),
                       ...){
  ## this function plots the results of an ASR on a given tree
  require(ape)
  # checks which package has been used to carry the ASR
  if(!inherits(x, "asr"))
    stop("x must be an object of class 'asr'")
  if(inherits(x$asr, "rerootingMethod")){
    lik <- x$asr$marginal.anc
  } else if(inherits(x$asr, "mpr")){
    lik <- x$asr$lik
  } else {
    stop("the format of likelihood scores is not recognized")}
  # applies the colour palette to plotted characters
  palette(palette)
  char <- x$data
  levels(char) <- palette()
  # plots the tree and adds a the tip and the node pies
  par(mar = rep(1, 4))
  plot(keep.tip(phy, x$tree$tip.label), 
       main = n, 
       cex = .8, 
       label.offset = 1,
       node.depth = 4,
       y.lim = y.lim, 
       edge.color = "gray50",
       edge.width = 1.2, 
       ...)
  tiplabels(pch = 21, col = tip.col, bg = char, cex = tip.size)
  nodelabels(pie = lik, col = pie.col, piecol = sort(unique(char)), cex = pie.size)
  # plots the phylogenetic signal values
  if(signal){
    sig <- round(phylosig.asr(x), 3)
    mtext(bquote(lambda==.(sig[1])), side = 1, line = -3, at = 1, adj = 0, cex = .8)
    mtext(bquote(p==.(sig[2])), side = 1, line = -2, at = 1, adj = 0, cex = .8)
  }
}

"pic.asr" <- function(obj, tree){
  require(ggplot2)
  require(ape)
  ## this function tests variable independence using phylogenetic contrasts
  if(!inherits(obj, "list"))
    stop("the object in input is not a list")
  # creation of indexes for all relevant pairs of variables
  m <- expand.grid(i = 1:length(obj), j = 1:length(obj))
  m <- m[m$j > m$i,]
  m <- m[order(m$i),]
  res <- c()
  for (n in 1:nrow(m)){
    i <- m$i[n]
    j <- m$j[n]
    varX <- obj[[i]]
    varY <- obj[[j]]
    if(!inherits(varX, "asr") | !inherits(varY, "asr"))
      stop("the objects must be of class 'asr'")
    X <- varX$data
    Y <- varY$data
    tips <- intersect(names(X), names(Y))
    phy <- keep.tip(tree, tips)
    picX <- pic(X[tips], phy)
    picY <- pic(Y[tips], phy)
    pic <- data.frame(picX, picY)
    colnames(pic) <- c("x", "y")
    mod <- lm(picY ~ picX - 1)
    plot <- ggplot(pic, aes(x, y)) +
      geom_point(shape = 1) +
      geom_smooth(method = "lm", formula = y ~ x - 1) +
      xlab(names(obj)[i]) +
      ylab(names(obj)[j])
    res[[n]] <- list(pic = pic, mod = summary(mod, correlation = T, signif.stars = T), plot = plot)
    names(res)[n] <- paste(names(obj)[j], "~", names(obj)[i])
  }
  class(res) <- "pic"
  return(res)
}

"pic.test" <- function(var){
  require(gtools)
  ## this function calculates phylogenetic independent contrasts for each variable
  test <- sapply(var, function(x){
    coeff <- data.frame(round(x$mod$coefficients, 3))
    if (nrow(coeff) == 0)
      coeff <- as.data.frame(matrix(NaN, 1, 4, dimnames = list(NULL, colnames(coeff))))
    coeff[, 5]  <- round(x$mod$r.squared, 3)
    return(coeff)
  })
  res <- data.frame(apply(test, 1, as.numeric), row.names = colnames(test))
  colnames(res) <- c("Estimate", "Std. Error", "t value", "Pr(>|t|)", "r.squared")
  res$VarX <- substr(rownames(res), regexpr("~ ", rownames(res))+2, nchar(rownames(res)))
  res$VarY <- substr(rownames(res), 1, regexpr(" ~", rownames(res)) - 1)
  rownames(res) <- NULL
  res <- res[, c(6:7, 1:5)]
  res$' ' <- stars.pval(res$'Pr(>|t|)')
  return(res)
}


##>>  Time calibration of tree <<##

tree <- read.tree("tree.tre")
taxa <- tree$tip.label
ages <- read.csv("ages.csv", sep = , header = T, row.names = 1) #ages dowloaded from the PbDb
caltree <- timePaleoPhy(tree, ages, type = "equal", vartime = 1)
plot(caltree, cex = .6)


##>>  Ancestral State Reconstruction  <<##

#>  Data import  <#

var <- c("pco", "nao", "naf", "ncf")
loci <- read.csv("loci.csv", sep = ";", row.name = 1, header = T, na.strings = "?")
loci <- lapply(var, function(x){
  df <- loci[taxa, grepl(x, names(loci))]
  lapply(df, function(x) factor(x, levels = 0:1))
  colnames(df) <- c("C", "T", "L", "S", "Ca")
  return(df)
})
names(loci) <- var

#>  Maximum likelihood  <#

asML <- lapply(loci, function(x){
  multi <- sapply(x, function(var) length(unique(var, na.rm = T))) > 2
  subset <- x[, multi]
  lapply(subset, function(var) asr(var, taxa = taxa, phy = caltree, model = "SYM"))
})

#>  Parsimony  <#

asPM <- lapply(loci, function(x){                                                        
  multi <- sapply(x, function(var) length(unique(var, na.rm = T))) > 2
  subset <- x[, multi]
  lapply(subset, function(var) mpr(var, taxa = taxa, phy = caltree, mod = "sequential"))
})

##>>  Plots  <<##

old.par <- par()

#> Maximum likelihood <#

mapply(x = asML, n = names(asML), function(x, n){
  nrow <- ceiling(length(x) / 3)
  if(nrow == 1) y.lim <- c(-40,40) else y.lim <- NULL
  par(mfrow = c(nrow,3))
  for(i in 1:length(x)){
    main <- paste(n, "-", names(x)[i])
    plot.asr(x[[i]], n = main, phy = tree, y.lim = y.lim)
  }
})

#> Parsimony <#

par(old.par)
mapply(x = asPM, n = names(asPM), function(x, n){
  nrow <- ceiling(length(x) / 3)W
  if(nrow == 1) y.lim <- c(-40,40) else y.lim <- NULL
  par(mfrow = c(nrow,3))
  for(i in 1:length(x)){
    main <- paste(n, "-", names(x)[i])
    plot.asr(x[[i]], n = main, phy = tree, y.lim = y.lim)
  }
})
```
